# Evidence of adoption, monozygotic twinning, and low inbreeding rates in a large genetic pedigree of polar bears

**DOI:** 10.1101/034009

**Authors:** René M. Malenfant, David W. Coltman, Evan S. Richardson, Nicholas J. Lunn, Ian Stirling, Elizabeth Adamowicz, Corey S. Davis

## Abstract

Multigenerational pedigrees have been developed for free-ranging populations of many species, are frequently used to describe mating systems, and are used in studies of quantitative genetics. Here, we document the development of a 4449-individual pedigree for the Western Hudson Bay subpopulation of polar bears *(Ursus maritimus)*, created from relationships inferred from field and genetic data collected over six generations of bears sampled between 1966 and 2011. Microsatellite genotypes for 22–25 loci were obtained for 2945 individuals, and parentage analysis was performed using the program FRANz, including additional offspring–dam associations known only from capture data. Parentage assignments for a subset of 859 individuals were confirmed using an independent high-density set of single nucleotide polymorphisms. To account for unsampled males in our population, we performed half-sib–full-sib analysis to reconstruct males using the program COLONY, resulting in a final pedigree containing 2957 assigned maternities and 1861 assigned paternities with only one observed case of inbreeding between close relatives. During genotyping, we identified two independently captured two-year-old males with identical genotypes at all 25 loci, showing—for the first time—a case of monozygotic twinning among polar bears. In addition, we documented six new cases of cub adoption, which we attribute to cub misidentification or misdirected maternal care by a female bereaved of her young. Importantly, none of these adoptions could be attributed to reduced female vigilance caused by immobilization to facilitate scientific handling, as has previously been suggested.

## Introduction

Multigenerational pedigrees are useful in studies describing mating systems and for quantitative genetics research (Pemberton 2008) and large pedigrees have been developed for wild populations of many species, including red deer (*Cervus elaphus*; Slate et al. 2002), bighorn sheep (*Ovis canadensis*; Poissant et al. 2010), song sparrows *(Melospiza melodía*; Reid et al. 2011), and American red squirrels (*Tamiasciurus hudsonicus*; Taylor et al. 2012). However, because of the great effort and expense of sampling large carnivores, few pedigrees have been developed for ursid species, with most parentage analyses containing no more than a few hundred individuals, which are typically sampled non-invasively (Cronin et al. 2005; De Barba et al. 2010; Itoh et al. 2012; Onorato et al. 2004; however, cf. Proctor et al. 2004; Bellemain et al. 2006). To date, the largest parentage analysis of polar bears *(Ursus maritimus)* was based on 583 individuals in the Barents Sea (Zeyl et al. 2009a; Zeyl et al. 2009b), which showed that polar bears exhibit serial monogamy, male-biased dispersal, and that inbreeding between close relatives is rare.

Polar bears are large carnivores that occur at low densities throughout the circumpolar Arctic and subarctic regions. They have a polygynous mating system (Derocher et al. 2010), typically breeding between late March and June, with females giving birth to 1–3 cubs in November–December while overwintering in maternity dens. Females emerge from dens in early spring, and are the only providers of parental care until their cubs become independent—typically at about 2.5 years old (Ramsay and Stirling 1988). Though family groups tend to avoid other bears—perhaps to avoid cannibalism and other conspecific aggression (Taylor et al. 1985)— cases of adoption have previously been documented (Atkinson et al. 1996; Belikov 1976; Derocher and Wiig 1999; Lunn et al. 2000; Saunders 2005; Vibe 1976). Although adoption has been known to occur in the Barents Sea subpopulation (Derocher and Wiig 1999), Zeyl et al. (2009a, 2009b) did not report any cases of adoption, perhaps because of the infrequency of occurrence, combined with the study’s comparatively small sample size.

Adoption has been observed in more than 60 mammalian species (Gorrell et al. 2010), and its occurrence requires special explanation due to the extremely high cost of milk provision to adopted young (e.g., Clutton-Brock et al. 1989). Allonursing and adoption may be explained adaptively through kin selection, reciprocal altruism, evacuation of excess milk, or through a gain in parenting experience (Roulin 2002). Alternately, adoption and allonursing may simply be the result of error, occurring especially when a reproductive individual is already hormonally or behaviourally primed to provide parental care and is bereaved of their young (Riedman 1982). Most empirical studies support the kin selection, milk evacuation, or misdirected parental care hypotheses (Roulin 2002). Amongst polar bears, adoption has been attributed to misdirected parental care caused by cub misidentification (Lunn et al. 2000), which may be caused by confusion due to the immobilization of adult females, which is necessary for scientific handling (Derocher and Wiig 1999).

Like adoption, monozygotic twinning is taxonomically widespread but infrequent, and although well described in humans (e.g., Bulmer 1970) and cattle (e.g., Silva del Río et al. 2006), few cases have been documented in wildlife species. Monozygotic quadruplets are the normal mode of reproduction among some species of armadillos (Hardy 1995), and monozygotic twins have been identified in lesser flat-headed bats *(Tylonycteris pachypus*; Hua et al. 2011), in wolves *(Canis lupus*; Carmichael et al. 2009), among some species of pinnipeds (Spotte 1982), including Antarctic fur seals *(Arctocephalus gazella*; Hoffman and Forcada 2009), and possibly in mule deer (Anderson and Wallmo 1984). The apparent scarcity of monozygotic twins is partially attributable to the difficulty of identifying them, as this requires genetic or embryological confirmation. For instance, conjoined twins, which develop from monozygotic twins and are therefore far rarer, are phenotypically conspicuous, and at least 20 cases of conjoined twinning in wildlife species have been published (Kompanje and Hermans 2008). To our knowledge, identical twinning (or conjoined twinning) has never been detected in any species of bear, although no studies have had large enough sample sizes to reliably detect such a rare event.

In this paper, we present a large pedigree of polar bears comprising 4449 individuals from the Western Hudson Bay subpopulation captured over six bear generations in northeastern Manitoba, Canada between 1966 and 2011. We document six new cases of cub adoption, and show—for the first time—an instance of monozygotic twinning among polar bears. Further, we find no cases of inbreeding between first-degree relatives. This pedigree is now being used to determine the mating system of polar bears (Richardson 2014), and in future studies, this pedigree will be used to determine the heritabilities of various body size metrics, some of which have been declining in this subpopulation for decades (Stirling and Derocher 2012).

## Methods

### Sample collection

Most tissue samples were collected from bears that were immobilized and handled as part of long-term ecological studies of polar bears in Western Hudson Bay. However, a small number of samples were collected from bears captured by Manitoba Conservation staff near the community of Churchill as part of the Polar Bear Alert Program (Kearney 1989) or from polar bears harvested each year as part of a legal, regulated subsistence hunt by Inuit living along the coast of western Hudson Bay (Derocher et al. 1997; Taylor et al. 2008). Sampling locations are shown in Figure 1. During the first handling of each captured individual, he/she was assigned a unique ID applied as a permanent tattoo on the inside of the upper lip and affixed as a plastic tag in each ear. Skin samples were collected by retaining leftover pinnal tissue from ear-tagging or from adipose tissue samples collected using a 6-mm biopsy punch of superficial fat on the rump (Ramsay et al. 1992; Thiemann et al. 2008). A temporary paint mark is also applied to avoid recapture of the same individual twice in one season. Blood samples were collected by drawing blood from a femoral vein into a sterile vacutainer. All samples were stored at −80°C until DNA extraction. If the age of a newly sampled individual was unknown (i.e., not a cub-of-the-year or a dependent yearling), a vestigial premolar tooth was extracted for age determination using measurement of cementum annulus deposition (Calvert and Ramsay 1998). All individuals handled by Environment Canada were sampled in autumn and were selected for handling indiscriminately of age or sex; however, in every year from 1980 onward (except for 1985 and 1986), a springtime sampling effort was also included, in which only adult females and their cubs-of-the-year—which had recently emerged from maternity dens—were handled.

**Figure 1.**
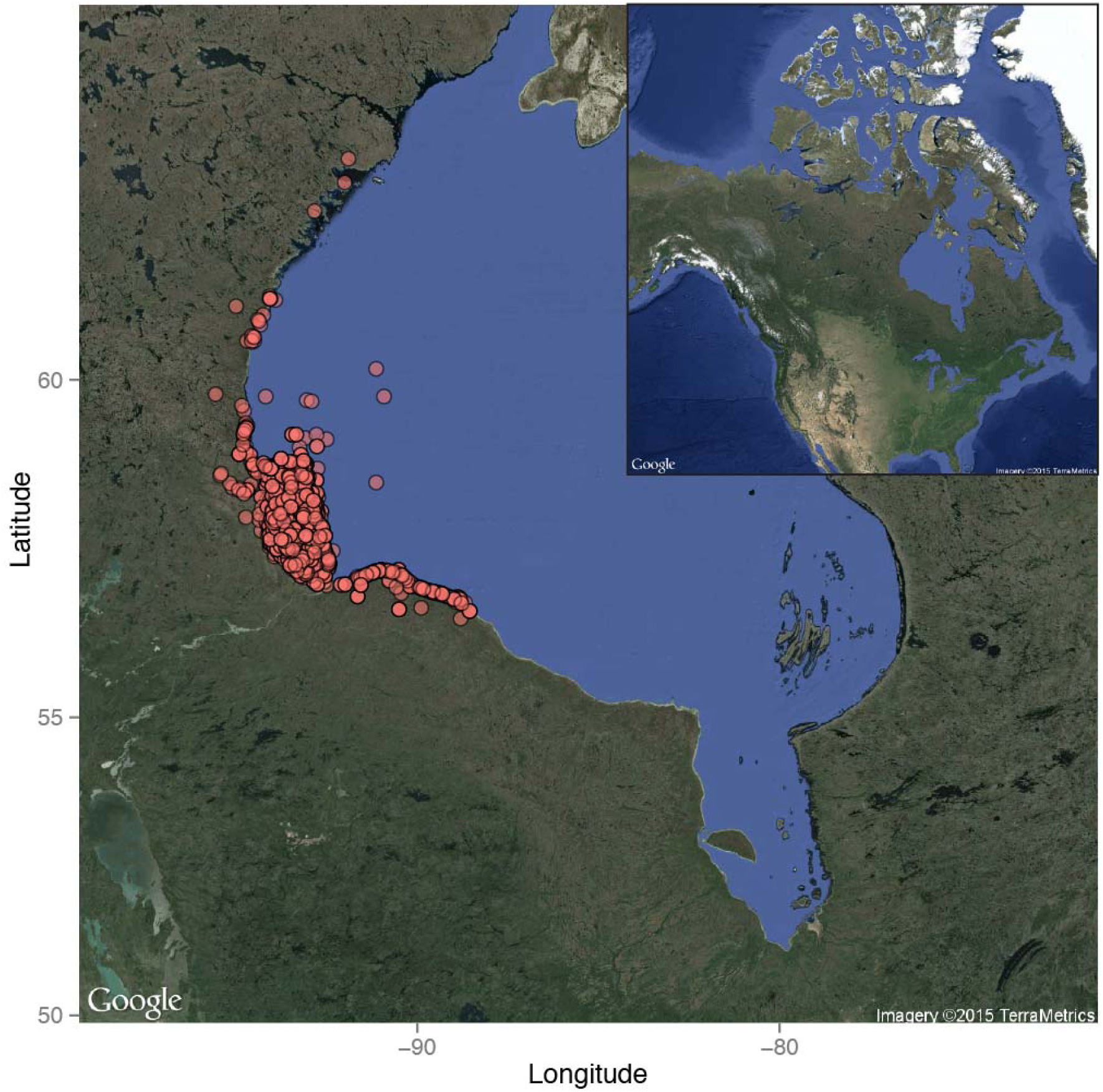
Sampling locations of bears included in the Western Hudson Bay pedigree. Imagery ©2015 TerraMetrics, Inc. (www.terrametrics.com), accessed via Google Maps and ggmap 2.4 (Kahle and Wickham 2013).

### DNA extraction and microsatellite genotyping

Total genomic DNA was extracted from fat, skin, or leukocytes recovered from ACK-lysed blood using DNeasy Blood & Tissue Kits (Qiagen, Hilden, Germany). We genotyped 2945 individuals born between 1960 and 2011, including duplicate samples from 69 individuals included to estimate genotyping error rates. Individuals born before 2006 were genotyped at all 25 microsatellite loci (Table S1, Supplementary Material); however, because of changes to the genotyping protocol made in 2012 to streamline microsatellite multiplexing, individuals born from 2006 onward were genotyped at 24 loci (excluding CXX173). PCR products from microsatellite amplifications were resolved on an Applied Biosystems 377 DNA Sequencer, 3100-Avant DNA Analyzer, or a 3730 DNA Analyzer, and sized relative to Genescan size standards. Genotyping was performed using the programs Genotyper and Genemapper (Applied Biosystems, Foster City, CA, USA).

### Genetic diversity, tests of disequilibrium, and statistics

The number of observed alleles (*N_A_*), observed heterozygosities *(H_O_)*, expected heterozygosities *(H_E_)*, probabilities of exclusion (*P_ex_*), and probabilities of identity (*P_ID_*) were calculated using GenAlex 6.5 (Peakall and Smouse 2006; Peakall and Smouse 2012); *P_ex_* was calculated using the formula for single unknown parent exclusion from Jamieson and Taylor (1997). Departures from Hardy–Weinberg equilibrium (HWE) and linkage disequilibrium (LD) were assessed with exact tests (Guo and Thompson 1992) and a Markov chain (dememorization number = 5000, number of batches = 1000, number of iterations per batch = 2000) using Genepop on the Web 4.2 (Raymond and Rousset 1995; Rousset 2008). The total genotyping error rate (calculated as the sum of the allelic dropout rate (E1) and false allele rate (E2), Table S1) was calculated from duplicate samples using the program Pedant 1.0 (Johnson and Haydon 2007) using 100,000 replicates. Unless otherwise indicated, all other statistics were calculated in R 3.1.3 (R Core Team 2015), and a significance level of *α*=0.05 was used for all tests.

### Pedigree generation and adoption detection

We used the program FRANz 2.0 (Riester et al. 2009; Riester et al. 2010) to generate an initial pedigree from our microsatellite data, using a sub-pedigree comprising known mother–cub relations from field data, individuals’ years of birth (and death, if known) and the settings specified in Table 1. FRANz generates a multigenerational pedigree in a single step, using simulations to determine expected parent–offspring mismatch rates, simulated annealing to estimate the maximum-likelihood pedigree, and Metropolis-coupled Markov chain Monte Carlo sampling to calculate parentage posterior probabilities. As part of its simulation step, FRANz uses the empirical distribution of parent–offspring mismatch rates to identify problematic parental assignments in the sub-pedigree (Figure 2), which we have classified here as adoptions. Although our field records contain no instances of females younger than four years old (or older than 28 years old) having successfully given birth, we specified a possible reproductive age range of 2–32 (at time of parturition) to account for outliers amongst unobserved maternities as well as possible errors in tooth aging. For simplicity, we also used this same age range for males; males generally do not become reproductively active until at least their fifth or sixth year, though they may begin to produce spermatozoa at age two (Rosing-Asvid et al. 2002). The oldest female and male polar bears captured from the Western Hudson Bay subpopulation were 32 and 29 years old, respectively.

**Figure 2.**
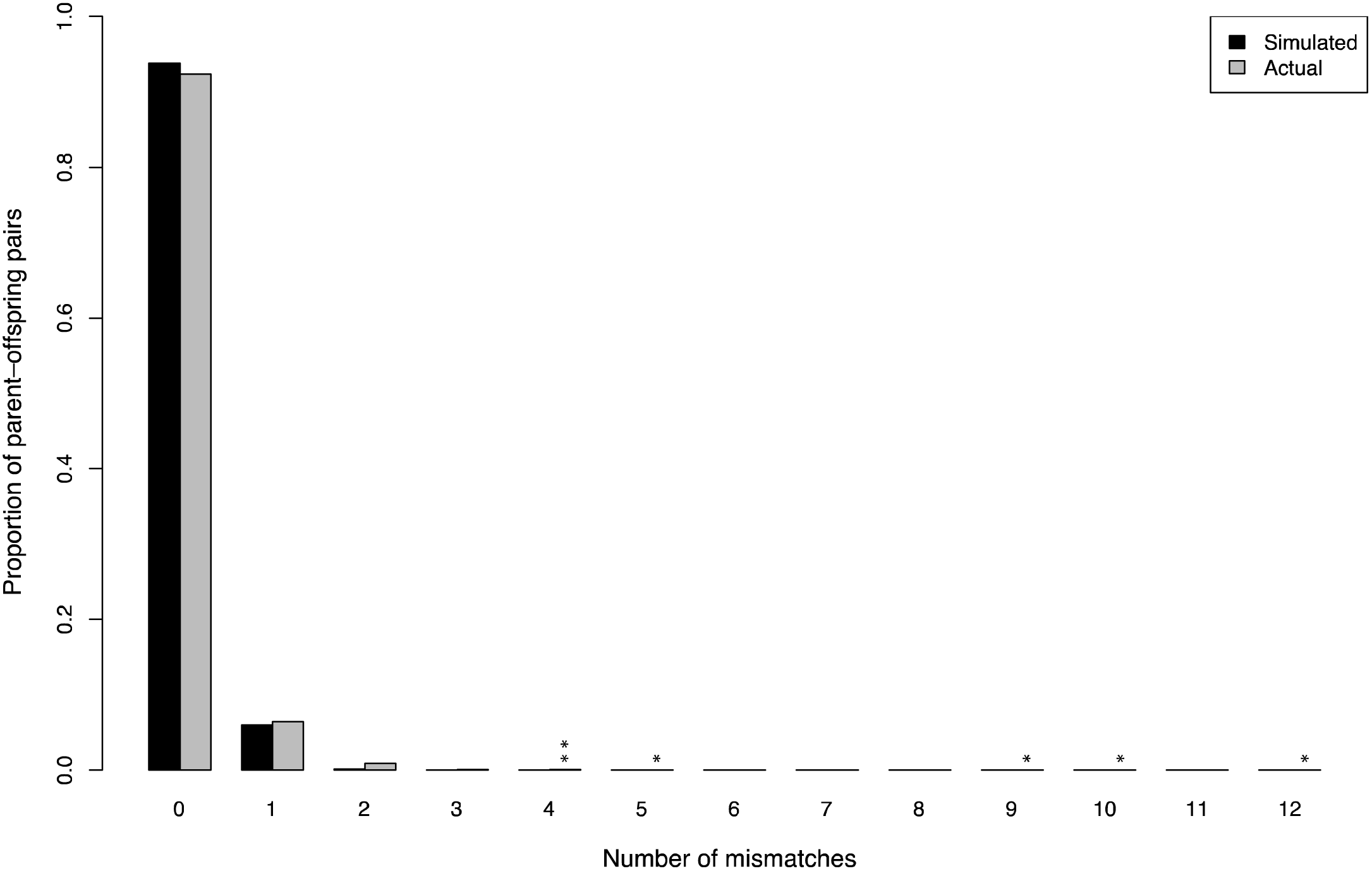
Number of mismatched microsatellite loci for simulated and actual parentage assignments. Simulation results represent 2,000,000 parent–offspring pairs generated in FRANz using empirically estimated error rates. Asterisks indicate putatively adopted individuals.

**Table 1.** Non-default settings used for pedigree generation in the program FRANz.

| Parameter | Setting |
| --- | --- |
| Female reproductive age range | 2–32 |
| Male reproductive age range | 2–32 |
| Maximum number of candidate mothers | 5,000 |
| Maximum number of candidate fathers | 5,000 |
| Number of simulation iterations | 1,000,000 |
| Maximum number of simulated annealing iterations | 1,000,000,000 |
| Number of Metropolis–Hastings burn-in iterations | 5,000,000 |
| Number of Metropolis–Hastings iterations | 30,000,000 |
| Metropolis–Hastings sampling frequency | 100 |

To detect the genetic mothers of adopted individuals, we removed their links from our field-data sub-pedigree and re-ran FRANz. Then, to validate and error-correct the resultant pedigree, we used the program VIPER 1.01 (Paterson et al. 2012) to examine the inheritance of 4,475 single-nucleotide polymorphisms (SNPs) genotyped in a 859-individual subset of pedigreed bears. These SNPs were developed from transcriptomic and RAD sequencing and were genotyped with high fidelity in all individuals using a recently developed 9K Illumina BeadChip for polar bears (Malenfant et al. 2015). We removed all pedigree links displaying more than one SNP inheritance error, which we determined as a cutoff based on the empirical distribution of inheritance errors.

Finally, to account for a lower proportion of males than females being sampled in our data (which led to proportionally fewer paternity than maternity assignments), we used the program COLONY 2.0 (Jones and Wang 2010) to generate hypothetical sires and differentiate between full siblings and maternal half siblings. To reduce false paternity assignments, we limited candidate offspring in this analysis to 760 individuals having unassigned sires but genetically assigned dams. All individuals were pooled in a single analysis irrespective of birth year to allow for the possibility that a hypothetical male had sired multiple offspring across years. We allowed for male and female polygamy, using maternal and paternal sibship priors of 3.655 and 2.968 respectively, which were determined empirically from the pedigree. All pedigree statistics were calculated using the package pedantics 1.5 (Morrissey and Wilson 2010).

### Genetic relatedness

Asocial animals such as polar bears might choose to adopt nearby orphans if they are genetically related, as this can provide an inclusive fitness advantage to the foster parent (e.g., Gorrell et al. 2010). To determine if adopted cubs were genetically related to their foster mothers, we used the program COANCESTRY 1.0.1.2 (Wang 2011) to obtain the Queller-and-Goodnight (1989), Lynch-and-Ritland (1999), and Wang (2002) relatedness metrics using allele frequencies and error rates estimated from the full microsatellite dataset. Because all estimators gave similar results, only the Queller-and-Goodnight estimator results are presented. This estimator was designed for studies of kin selection and has the property that unrelated individuals are expected, on average, to have a relatedness of zero.

## Results

### Microsatellite genotypes

Complete 25-locus genotypes were obtained for 2418 individuals, 24-locus genotypes were obtained for 478 individuals, 23-locus genotypes were obtained for 34 individuals, and 22-locus genotypes were obtained for 15 individuals. The mean number of observed alleles (*N_A_*) was 7.6 (range: 3–10), and mean observed (*H_O_*) and expected *(He)* heterozygosities were both 0.672 (ranges: 0.112–0.847 and 0.112–0.840, respectively). Two loci, G1A and G10L, deviated from HWE *(P_G1A_* = 0.0028, *P_G1OL_* = 0.0064) but were not significantly out of HWE following a strict Bonferroni correction for multiple tests *(α_corrected_* = 0.002). Complete summary statistics for microsatellite loci are presented in Table S1. Thirty pairwise tests of LD were significant following strict Bonferroni correction, however, this was likely because our dataset contained many groups of related individuals. Combined probability of exclusion *(P_ex_)* over all loci is 0.99991; this drops to 0.99988 if CXX173 is excluded. Combined probability of identity (*P_ID_*) was 7.102×10^−23^ for unrelated individuals and 1.562×10^−9^ for full siblings; these increase to 5.020×10^−22^ and 3.551×10^−9^ respectively if CXX173 is excluded. Total genotyping error rate was estimated at 0.36%. Fourteen successfully genotyped individuals were removed prior to pedigree generation because their year of birth was unknown.

### Pedigree statistics and inbreeding

We supplemented the remaining 2931 genotyped-and-aged individuals with 1225 individuals known only from field observation. FRANz assigned 2972 maternities using field and/or genetic data and 1105 paternities using genetic data alone. Based on SNP inheritance errors identified in VIPER, 4 offspring–sire links out of 163 (≈2.5%) and 15 offspring–dam links out of 465 (≈3.2%) were removed. In all 15 of these cases, offspring–dam relationships had been inferred using genetic data only (i.e., they were not based on field observations). COLONY reconstructed 293 sires, which collectively accounted for 760 paternal assignments (mean ± SD offspring per reconstructed sire = 2.6 ± 1.4), and brought the pedigree to 4449 individuals in total. Nine females aged 3 years or younger and males aged 2 years or younger (at time of conception) were assigned as parents in the final version of the pedigree, though the ages of 17 of these 18 individuals were uncertain as they were derived from tooth-aging estimates. The oldest dam and sire assigned in the pedigree were 28 and 30 respectively at the time of conception.

Including COLONY-reconstructed individuals, the pedigree contains 1381 founders (i.e., individuals of unknown parentage) and extends to six generations for some individuals. Of 382 mating events in which the identities of both parents and at least one grandparent on each side were known (a necessity for detecting inbreeding), only three individuals had non-zero inbreeding coefficients: X11088, X11089, and X11389. X11088 and X11089 are littermates born in 1989 to X10668 after mating with her half-brother X10497; X11389 was supposedly born to X09396 and her brother X09398 (however, cf. the Discussion regarding this mating). A graphical view of the pedigree and complete pedigree statistics and are given in Figure 3 and Table S2 of the Supplementary Material, respectively.

**Figure 3.**
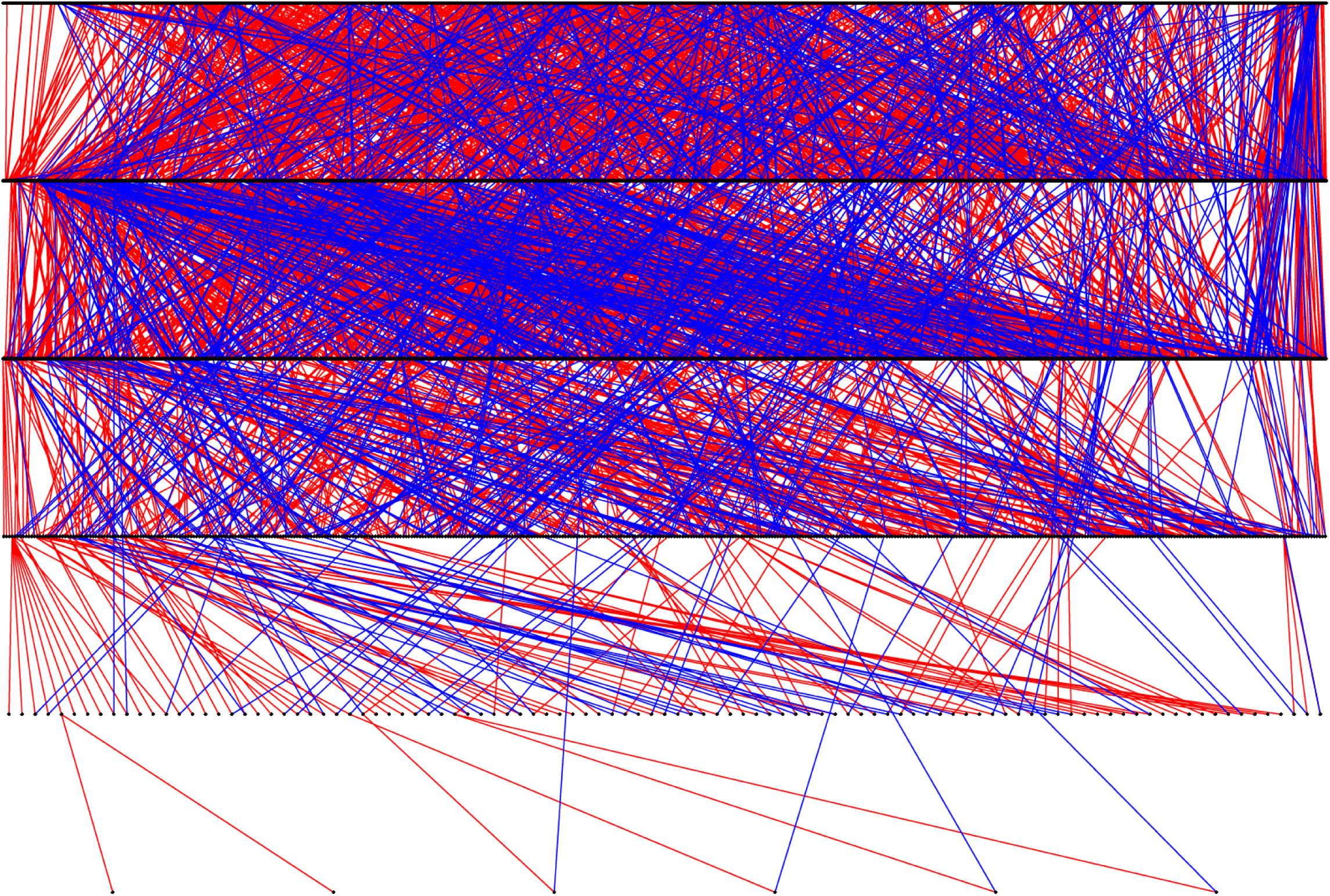
Graphical representation of the 4449-individual polar bear pedigree described in this paper. Each point is an individual bear. Maternities are represented by red lines; paternities are represented by blue lines.

### Monozygotic twinning

We detected one pair of identical twins among 574 genotyped twin litters and 37 genotyped triplet litters: cubs X17324 and X17326 match at all 25 loci (Supplementary Material 2). Both individuals were independent two-year olds at the time of capture (November 10 and 11, 2003, respectively), and were handled ~3.5 km apart on opposite sides of the Churchill airport. They were known not to have been recaptures of the same individual because the second-captured individual (X17326) lacked the temporary paint mark and did not have a permanent tattoo or ear tags. If dizygotic, the probability of these cubs sharing a genotype at all 25 loci is 1.64×10^−11^, as calculated from the full genotypes of both parents. To discount the possibility that these identical genotypes were the result of sample mix-up, we reconfirmed the genotype of X17324 using a second, independently collected tissue sample; unfortunately, a second sample for X17326 was not available. However, because these individuals were handled on different days, the probability of sample mix-up during fieldwork is extremely low.

### Cases of adoption

We identified six previously undetected cases of adoption occurring between 1981 and 2004 and identified four of the six genetic mothers (Table 2, Supplementary Material S2). In five of these cases, cubs were adopted during their first year of life; in the remaining case (X09059), it was unclear if the cub was adopted during its first or second year. In two cases, adoptive mothers were observed to have fostered cubs for at least a year, and from later capture and harvest records, it is known that at least five of six adopted cubs survived to independence (though the fate of X11097 is unknown.) Although five of the six adopted cubs were female, there was no statistical evidence of preference to adopt females over males (binomial test of 1:1 ratio: *P* = 0.2188). All adopted cubs appeared to be unrelated to their adoptive mothers: average adoptee–adopter relatedness is −0.038, and 95% confidence intervals for the Queller-and-Goodnight relatedness estimators are not significantly different from 0 in all cases. In two of the six adoption cases, females were also accompanied by their own genetic offspring.

**Table 2.** New cases of polar bear adoption reported in this paper. *P_M_* = proportion of loci mismatched between cub and candidate mother; *r_QG_* = Queller-and-Goodnight relatedness between cub and the adoptive mother. In two cases (represented as “—”), the true dam could not be determined. Asterisks denote individuals whose genotypes could be confirmed by genetic assignment of their other observed offspring to them; most cubs’ genotypes could not be confirmed in this manner because they are not known to have parented offspring.

| Cub | Sex | Year of birth | Observed dam ($P_M$) | Date(s) observed together | Age of observed dam | Other cubs in litter | $r_{QG}$ (95% CI) | Inferred dam ($P_M$) | Survived to independence |
| --- | --- | --- | --- | --- | --- | --- | --- | --- | --- |
| X09059* | Female | 1980 | X05562*<br>(10/25) | Aug. 10, 1981 | 7 | 0 | −0.19<br>(−0.36 – 0.03) | X09913*<br>(0/25) | Yes |
| X10608 | Female | 1987 | X10607*<br>(5/25) | Sept. 24, 1987 | 8 | 0 | −0.05<br>(−0.22 – 0.12) | — | Yes |
| X11097 | Female | 1989 | X05668*<br>(9/25) | Mar. 22, 1989 | 6 | 2 | −0.07<br>(−0.35 – 0.14) | X11456*<br>(0/25) | Unknown |
| X17069 | Male | 1998 | X03318*<br>(4/25) | Nov. 30, 1998 | 24 | 0 | 0.17<br>(0.00 – 0.36) | — | Yes |
| X17294* | Female | 2003 | X17082*<br>(4/25) | Sept. 22, 2003 –<br>Nov. 8, 2004 | 7–8 | 1 | 0.03<br>(−0.12 – 0.21) | X10688*<br>(0/25) | Yes |
| X19939 | Female | 2004 | X11940*<br>(12/25) | Sept. 18, 2004 –<br>Sept. 18, 2005 | 11–12 | 0 | −0.12<br>(−0.38 – 0.10) | X12273*<br>(1/25) | Yes |

## Discussion

### Inbreeding

Active inbreeding avoidance is often presumed to be common amongst animals because of reduced fitness of inbred offspring (Keller and Waller 2002), though tolerance of—or even preference for—inbreeding may occur because of inclusive fitness benefits (Szulkin et al. 2013). When inbreeding avoidance does occur, it is generally attributed to mate choice or sex-biased dispersal (Pusey and Wolf 1996). However, sex-biased dispersal may also occur for reasons unrelated to inbreeding, such as sex differences in the benefits of retaining a productive territory or avoidance of intersexual competition (Moore and Ali 1984). Little is known about inbreeding in polar bears, and primary among the motivations for developing this pedigree was the characterization of inbreeding in this subpopulation (Richardson et al. 2006).

We detected only two instances of incestuous mating: one between half-siblings X10668 and X10497 (producing X11088 and X11089), and another putative case between full-siblings X09396 and X09398 (producing X11389). However, in this latter case, X09398 is almost certainly a false paternity assignment: X09396 is an ungenotyped dam that was assigned using only field data, causing X09398 to be incorrectly assigned as a father because of allele-sharing with his sister. Therefore, after excluding this case, inbreeding among close relatives appears to be extremely rare in the Western Hudson Bay subpopulation, occurring only once among 382 mating events in which it could have been observed. For comparison, in a study of the Barents Sea subpopulation, a single instance of father–daughter inbreeding was detected amongst 22 matings between parents of known identity (Zeyl et al. 2009a), suggesting that the rate of mating between first-degree relatives was ~4.5%.

Polar bears exhibit low variation at major histocompatibility loci (Weber et al. 2013), which are thought to play an important role in kin recognition (Villinger and Waldman 2012), and it has also been suggested that polar bears have undergone little selection for kin recognition because of low population densities (Lunn et al. 2000). If this is the case, then mate choice is unlikely to explain low inbreeding rates amongst polar bears. In contrast, our finding of little-to-no inbreeding in the Western Hudson Bay subpopulation may result from substantial dispersal and interbreeding between Western Hudson Bay and adjacent management units (Richardson 2014). Studies of American black bears *(Ursus americanus*; Costello et al. 2008) and brown bears *(U. arctos;* Bellemain et al. 2006) have found similar rates of close inbreeding, which were attributed to lack of opportunity resulting from low population density and male-biased dispersal. In another study, inbreeding avoidance was cited as the most likely cause of male natal dispersal among brown bears (Zedrosser et al. 2007). These findings are also likely to hold true for polar bears, which occur at even lower densities, and for which limited genetic evidence also suggests male-biased dispersal in some subpopulations (Zeyl et al. 2009b). However, because radio telemetry data for male polar bears is scarce, little is known about dispersal patterns in male polar bears and further study is needed.

### Monozygotic twinning

Inclusive fitness theory predicts the possible spread of genes for monozygotic twinning (Gleeson et al. 1994; Williams 1975), and though the reason for the rarity of monozygotic twinning is not well understood, it may be partially attributable to higher rates of spontaneous abortion for monozygotic twins (Livingston and Poland 1980) and lower survival of twins in species that normally bear only one offspring (e.g., Fricke 2001). Based on observed sex ratios of multi-cub litters, Ramsay and Stirling (1988) determined that monozygotic twinning was likely rare or absent among polar bears. Our study confirms that monozygotic polar bear twins are extremely rare, being found in less than 1/600 litters (≈0.17%) in our data. To our knowledge, this is the first confirmed record of monozygotic twinning among polar bears or any other ursid, and previous genetics studies of bears (e.g., Bellemain et al. 2006; Proctor et al. 2004; Zeyl et al. 2009a) likely failed to detect twins because of smaller sample sizes. Slightly higher rates of monozygotic twinning have been found for humans (0.35–0.4%; Bulmer 1970) and for cattle (0.33%; Silva del Río et al. 2006). In part, the lower estimate for polar bears may result from discounting the “invisible fraction” (Grafen 1988) of identical twins that were never observed because at least one cub died prior to emergence from the maternity den.

### Adoption

According to Hamilton’s (1964) theory of kin selection, natural selection will favour a heritable predisposition for altruistic behaviour when *C* < *rB*, where *C* is the fitness cost to the altruist, *B* is the fitness benefit to recipient, and *r* is the relatedness between these individuals. Thus, kin selection requires greater-than-average relatedness between altruist and recipient, and relatedness must be particularly high to account for such energetically costly behaviours as adoption and nursing. Lunn et al. (2000) ruled out kin selection as an explanation for three previous cases of polar bear adoption based on low genetic relatedness. Our results reinforce the finding that adopted cubs and their foster mothers are unrelated, and that kin selection does not appear to drive adoption in this population. Though reciprocal care of offspring has been observed in polar bears (Lunn 1986), given the polar bear’s low population density and generally asocial nature, reciprocal altruism is also extremely unlikely, and no reciprocal cases of adoption were observed in our data.

Milk evacuation may explain allonursing behaviour in some pinnipeds (Riedman and Boeuf 1982) and bats (Wilkinson 1992), however, it is extremely unlikely to explain adoption in polar bears. Whereas it is beneficial for pinnipeds and bats to be leaner to increase diving or flight efficiency (Roulin 2002), lean polar bears lack the energy storage and thermal benefits (Pond et al. 1992), as well as the reproductive benefits (Stirling et al. 1999) associated with body condition. This is particularly true in the Western Hudson Bay subpopulation, where mothers may fast for four or more months during the ice-free period each year. In this subpopulation, a female’s ability to maintain pregnancy (Derocher et al. 1992) and the survival of her own cubs (Derocher and Stirling 1996) are mass-dependent so that a female would gain no apparent benefit from milk evacuation. Since it appears that all six foster mothers had birthed genetic litters by the time of adoption, the parental experience hypothesis is also unlikely to account for any of these adoptions (Roulin 2002).

Adopted cubs were captured alone with their foster mother in four of six cases, and in all these cases, the adopted cub is known to have survived to independence, implying the provisioning of milk by the mother to the adoptive offspring, as has been observed directly in at least once instance of fostering (Belikov 1976). Because spontaneous lactation is not believed to occur in most species and has only been consistently demonstrated among dwarf mongooses *(Helogale parvula)* suckling close relatives (Creel et al. 1991), it is highly unlikely to explain allonursing of alien offspring among polar bears. This suggests that these cubs’ adoptions coincided with the loss of the females’ biological litters (either due to death or because of unintentional cub-swapping with another female), while females were biologically capable of suckling. In our remaining two adoption cases, cub misidentification is the most likely explanation, as adopted cubs were accompanied by the female’s own biological offspring. Cub mixing sometimes occurs in both polar bears and brown bears (Glenn et al. 1976; Lunn 1986), and in any of these adoption cases, cub-swapping may have occurred due to simple misidentification during periods of high bear density, such as springtime den emergence or the autumn fasting period ashore (Derocher and Stirling 1990; Ramsay and Stirling 1988). However, we note that in at least one previously observed case of adoption, it has been proposed that a female with two cubs of her own adopted two of another female’s cubs she was killed in a fight (Vibe 1976), and cubs may also become separated from their mothers if males drive them off in order to mate with her (I. Stirling, unpublished data).

It has been suggested that scientific handling may increase the probability of cub abandonment or adoption if maternal vigilance is reduced during the time it takes to fully recover from immobilization (Derocher and Wiig 1999). Importantly, we found no evidence to support this hypothesis. We were able to identify four of the six genetic mothers, two of which had not been captured for five years prior to the adoption, and the remaining two of which were not captured until after the adoption. Likewise, none of the six foster mothers was captured in the period between the adopted cub’s birth and their first observation together. This finding corresponds with a number of studies that have failed to find a significant negative correlation between scientific handling and litter size (Amstrup 1993; Lunn et al. 2004), cub survival (Ramsay and Stirling 1986; Rode et al. 2014), or on the cohesion of family groups (Messier 2000).

## Acknowledgments

The authors would like to thank the Manitoba Department of Conservation and the Government of Nunavut for providing some of the samples, as well as Dennis Andriashek and Wendy Calvert for long-term data collection and maintenance. We would also like to acknowledge the contributions of Andrew Derocher and the late Malcolm Ramsay, who gathered some of the field data used in this study. DNA extractions for most pre-2006 tissue samples were conducted by Jennifer Bonneville Davis. This project was funded by grants to CSD from Environment Canada and by a Natural Sciences and Engineering Research Council Discovery Grant to DWC (Grant ID 312207–2011). RMM is funded by scholarships from Alberta Innovates Technology Futures, the University of Alberta, and the Province of Alberta. Financial and logistical support of the long-term study of polar bears in western Hudson Bay have been provided by Care for the Wild International, the Churchill Northern Studies Centre, Environment Canada, the Isdell Family Foundation, Manitoba Conservation, Natural Sciences and Engineering Research Council, Nunavut Wildlife Research Trust Fund, Parks Canada Agency, the Strategic Technology Applications of Genomics in the Environment (STAGE) funding program, Wildlife Media Inc., World Wildlife Fund (Canada), and World Wildlife Fund Arctic Programme. Josh Miller and Jamie Gorrell provided advice and comments on an earlier version of the manuscript.

## Ethical standards

All applicable international, national, and/or institutional guidelines for the care and use of animals were followed. Environment Canada's animal-handling procedures were approved annually by their Prairie and Northern Region Animal Care Committee, and all research was conducted under wildlife research permits issued by the Province of Manitoba and by Parks Canada Agency.

## References

Amstrup SC (1993) Human disturbances of denning polar bears in Alaska. Arctic 46: 246–250.

Anderson AE, Wallmo OC (1984) Odocoileus hemionus. Mammalian Species 219: 1–9;.

Atkinson SN, Cattet MRL, Polischuk SC, Ramsay MA (1996) A case of offspring adoption in free-ranging polar bears *(Ursus maritimus)*. Arctic 49: 94–96.

Belikov SE (1976) Behavioral aspects of the polar bear, *Ursus maritimus*. Bears: Their Biology and Management 3:37–40. doi: 10.2307/3872752

Bellemain E, Zedrosser A, Manel S, Waits LP, Taberlet P, Swenson JE (2006) The dilemma of female mate selection in the brown bear, a species with sexually selected infanticide. P R Soc B 273:283–291. doi: 10.1098/Rspb.2005.3331

Bulmer MG (1970) The Biology of Twinning in Man. Clarendon Press, Oxford, UK Calvert W, Ramsay MA (1998) Evaluation of age determination of polar bears by counts of cementum growth layer groups. Ursus 10:449–453. doi: 10.2307/3873156

Carmichael L, Nagy JA, Strobeck C (2009) Monozygotic twin wolves with divergent life histories. Arctic 61:329–331. doi: 10.14430/arctic29 329-331

Clutton-Brock TH, Albon SD, Guinness FE (1989) Fitness costs of gestation and lactation in wild mammals. Nature 337: 260–262.

Costello CM, Creel SR, Kalinowski ST, Vu NV, Quigley HB (2008) Sex-biased natal dispersal and inbreeding avoidance in American black bears as revealed by spatial genetic analyses. Mol Ecol 17:4713–4723. doi: 10.1111/j.1365-294X.2008.03930.x

Creel SR, Monfort SL, Wildt DE, Waser PM (1991) Spontaneous lactation is an adaptive result of pseudopregnancy. Nature 351: 660–662.

Cronin MA, Shideler R, Waits L, Nelson RJ (2005) Genetic variation and relatedness on grizzly bears in the Prudhoe Bay region and adjacent areas in northern Alaska. Ursus 16:70–84. doi: 10.2192/1537-6176(2005)016[0070:Gvarig]2.0.Co;2

De Barba M, Waits LP, Garton EO, Genovesi P, Randi E, Mustoni A, Groff C (2010) The power of genetic monitoring for studying demography, ecology and genetics of a reintroduced brown bear population. Mol Ecol 19:3938–3951. doi: 10.1111/J.1365-294x.2010.04791.X

Derocher AE, Stirling I (1990) Distribution of polar bears *(Ursus maritimus)* during the ice-free period in western Hudson Bay. Can J Zool 68: 1395–1403.

Derocher AE, Stirling I, Andriashek D (1992) Pregnancy rates and serum progesterone levels of polar bears in western Hudson Bay. Can J Zool 70:561–566. doi: 10.1139/z92-084

Derocher AE, Stirling I (1996) Aspects of survival in juvenile polar bears. Can J Zool 74: 1246–1252.

Derocher AE, Stirling I, Calvert W (1997) Male-biased harvesting of polar bears in western Hudson Bay. J Wildl Manage 61: 1075–1082.

Derocher AE, Wiig Ø (1999) Observation of adoption in polar bears *(Ursus maritimus)*. Arctic 52: 413–415.

Derocher AE, Andersen M, Wiig Ø, Aars J (2010) Sexual dimorphism and the mating ecology of polar bears *(Ursus maritimus)* at Svalbard. Behav Ecol Sociobiol 64:939–946. doi: 10.1007/s00265-010-0909-0

Fricke PM (2001) Review: twinning in dairy cattle. The Professional Animal Scientist 17:61–67.

Gleeson SK, Clark AB, Dugatkin LA (1994) Monozygotic twinning: an evolutionary hypothesis. Proceedings of the National Academy of Sciences 91: 11363–11367.

Glenn LP, Lentfer JW, Faro JB, Miller LH (1976) Reproductive biology of female brown bears *(Ursus arctos)*, McNeil River, Alaska. Bears: Their Biology and Management 3:381–390. doi: 10.2307/3872788

Gorrell JC, McAdam AG, Coltman DW, Humphries MM, Boutin S (2010) Adopting kin enhances inclusive fitness in asocial red squirrels. Nat Commun 1:22. doi: 10.1038/Ncomms1022

Grafen A (1988) On the Uses of Data on Lifetime Reproductive Success. In: Clutton-Brock TH (ed) Reproductive Success. Univeristy of Chicago Press, Chicago, IL, pp 454–471

Guo SW, Thompson EA (1992) Performing the exact test of Hardy-Weinberg proportion for multiple alleles. Biometrics 48: 361–372.

Hamilton WD (1964) The genetical evolution of social behaviour. I. J Theor Biol 7:1–16. doi: 10.1016/0022-5193(64)90038-4

Hardy ICW (1995) Protagonists of polyembryony. Trends Ecol Evol 10:179–180. doi: http://dx.doi.org/10.1016/S0169-5347(00)89045-X

Hoffman JI, Forcada J (2009) Genetic analysis of twinning in Antarctic fur seals *(Arctocephalus gazella)*. J Mammal 90:621–628. doi: 10.1644/08-MAMM-A-264R1.1

Hua P, Zhang L, Zhu G, Jones G, Zhang S, Rossiter SJ (2011) Hierarchical polygyny in multiparous lesser flat-headed bats. Mol Ecol 20:3669–3680. doi: 10.1111/j.1365-294X.2011.05192.x

Itoh T, Sato Y, Kobayashi K, Mano T, Iwata R (2012) Effective dispersal of brown bears (Ursus arctos) in eastern Hokkaido, inferred from analyses of mitochondrial DNA and microsatellites. Mamm Study 37: 29–41.

Jamieson A, Taylor SS (1997) Comparisons of three probability formulae for parentage exclusion. Anim Genet 28:397–400. doi: 10.1111/J.1365-2052.1997.00186.X

Johnson PCD, Haydon DT (2007) Maximum-likelihood estimation of allelic dropout and false allele error rates from microsatellite genotypes in the absence of reference data. Genetics 175:827–842. doi: 10.1534/Genetics.106.064618

Jones OR, Wang J (2010) COLONY: a program for parentage and sibship inference from multilocus genotype data. Mol Ecol Resour 10:551–555. doi: 10.1111/J.1755-0998.2009.02787.X

Kahle D, Wickham H (2013) ggmap: spatial visualization with ggplot2. The R Journal 5: 144–161.

Kearney SR (1989) The Polar Bear Alert Program at Churchill, Manitoba. In: Bromley M (ed) Bear–People Conflicts: Proceedings of a Symposium on Management Strategies. Northwest Territories Department of Renewable Resources, Yellowknife, Northwest Territories, Canada, pp 83–92

Keller LF, Waller DM (2002) Inbreeding effects in wild populations. Trends Ecol Evol 17:230–241. doi: http://dx.doi.org/10.1016/S0169-5347(02)02489-8

Kompanje EJO, Hermans JJ (2008) Cephalopagus conjoined twins in a leopard cat *(Prionailurus bengalensis)*. J Wildl Dis 44:177–180. doi: 10.7589/0090-3558-44.1.177

Livingston JE, Poland BJ (1980) A study of spontaneoulsy aborted twins. Teratology 21:139–148. doi: 10.1002/tera.1420210202

Lunn NJ (1986) Observations of nonaggressive behavior between polar bear family groups. Can J Zool 64:2035–2037. doi: 10.1139/Z86-307

Lunn NJ, Paetkau D, Calvert W, Atkinson S, Taylor M, Strobeck C (2000) Cub adoption by polar bears *(Ursus maritimus):* determining relatedness with microsatellite markers. J Zool 251: 23–30.

Lunn NJ, Stirling I, Andriashek D, Richardson E (2004) Selection of maternity dens by female polar bears in western Hudson Bay, Canada and the effects of human disturbance. Polar Biol 27:350–356. doi: 10.1007/s00300-004-0604-6

Lynch M, Ritland K (1999) Estimation of pairwise relatedness with molecular markers. Genetics 152: 1753–1766.

Malenfant RM, Coltman DW, Davis CS (2015) Design of a 9K Illumina BeadChip for polar bears *(Ursus maritimus)* from RAD and transcriptome sequencing. Mol Ecol Resour 15:587–600. doi: 10.1111/1755-0998.12327

Messier F (2000) Effects of capturing, tagging and radio-collaring polar bears for research and management purposes in Nunavut and Northwest Territories. Department of Biology, University of Saskatchewan, Saskatoon, SK, Canada, pp 64

Moore J, Ali R (1984) Are dispersal and inbreeding avoidance related? Anim Behav 32:94–112. doi: http://dx.doi.org/10.1016/S0003-3472(84)80328-0

Morrissey MB, Wilson AJ (2010) PEDANTICS: an R package for pedigree-based genetic simulation and pedigree manipulation, characterization and viewing. Mol Ecol Resour 10:711–719. doi: 10.1111/J.1755-0998.2009.02817.X

Onorato DP, Hellgren EC, Van Den Bussche RA, Skiles JR (2004) Paternity and relatedness of American black bears recolonizing a desert montane island. Can J Zool 82:1201–1210. doi: 10.1139/Z04-097

Paterson T, Graham M, Kennedy J, Law A (2012) VIPER: a visualisation tool for exploring inheritance inconsistencies in genotyped pedigrees. BMC Bioinformatics 13:16. doi: 10.1186/1471-2105-13-s8-s5

Peakall R, Smouse PE (2006) GENALEX 6: genetic analysis in Excel. Population genetic software for teaching and research. Mol Ecol Notes 6:288–295. doi: 10.1111/J.1471-8286.2005.01155.X

Peakall R, Smouse PE (2012) GenAlEx 6.5: genetic analysis in Excel. Population genetic software for teaching and research-an update. Bioinformatics 28:2537–2539. doi: 10.1093/Bioinformatics/Bts460

Pemberton JM (2008) Wild pedigrees: the way forward. P R Soc B 275:613–621. doi: 10.1098/rspb.2007.1531

Poissant J, Hogg JT, Davis CS, Miller JM, Maddox JF, Coltman DW (2010) Genetic linkage map of a wild genome: genomic structure, recombination and sexual dimorphism in bighorn sheep. BMC Genomics 11. doi: 10.1186/1471-2164-11-524

Pond CM, Mattacks CA, Colby RH, Ramsay MA (1992) The anatomy, chemical composition, and metabolism of adipose tissue in wild polar bears *(Ursus maritimus)*. Can J Zool 70: 326–341.

Proctor MF, McLellan BN, Strobeck C, Barclay RMR (2004) Gender-specific dispersal distances of grizzly bears estimated by genetic analysis. Can J Zool 82:1108–1118. doi: 10.1139/Z04-077

Pusey A, Wolf M (1996) Inbreeding avoidance in animals. Trends Ecol Evol 11:201–206. doi: http://dx.doi.org/10.1016/0169-5347r96i10028-8

Queller DC, Goodnight KF (1989) Estimating relatedness using genetic markers. Evolution 43:258–275. doi: 10.2307/2409206

R Core Team (2015) R: A Language and Environment for Statistical Computing. R Foundation for Statistical Computing, Vienna, Austria,

Ramsay MA, Stirling I (1986) Long-term effects of drugging and handling free-ranging polar bears. The Journal of Wildlife Management 50:619–626. doi: 10.2307/3800972

Ramsay MA, Stirling I (1988) Reproductive biology and ecology of female polar bears *(Ursus maritimus)*. J Zool 214: 601–634.

Ramsay MA, Mattacks CA, Pond CM (1992) Seasonal and sex differences in the structure and chemical composition of adipose tissue in wild polar bears *(Ursus maritimus)*. J Zool 228: 533–544.

Raymond M, Rousset F (1995) GENEPOP (Version 1.2): population genetics software for exact tests and ecumenicism. J Hered 86: 248–249.

Reid JM, Arcese P, Sardell RJ, Keller LF (2011) Additive genetic variance, heritability, and inbreeding depression in male extra-pair reproductive success. The American Naturalist 177:177–187. doi: 10.1086/657977

Richardson E, Branigan M, Calvert W, Cattet M, Derocher AE, Doidge W, Hedman D, Lunn NJ, McLoughlin P, Obbard ME, Stirling I, Taylor M (2006) Research on polar bears in Canada 2001–2004. In: Aars J, Lunn NJ, Derocher AE (eds) Polar Bears: Proceedings of the 14th Working Meeting of the IUCN/SSC Polar Bear Specialist Group, 20–24 June 2005, Seattle, Washington, USA. IUCN, Gland, Switzerland,

Richardson ES (2014) The Mating System and Life History of the Polar Bear. University of Alberta

Riedman ML (1982) The evolution of alloparental care and adoption in mammals and birds. Q Rev Biol 57:405–435. doi: 10.1086/412936

Riedman ML, Boeuf BJ (1982) Mother-pup separation and adoption in northern elephant seals. Behav Ecol Sociobiol 11:203–215. doi: 10.1007/BF00300063

Riester M, Stadler PF, Klemm K (2009) FRANz: reconstruction of wild multi-generation pedigrees. Bioinformatics 25:2134–2139. doi: 10.1093/Bioinformatics/Btp064

Riester M, Stadler PF, Klemm K (2010) Reconstruction of pedigrees in clonal plant populations. Theor Popul Biol 78:109–117. doi: 10.1016/J.Tpb.2010.05.002

Rode KD, Pagano AM, Bromaghin JF, Atwood TC, Durner GM, Simac KS, Amstrup SC (2014) Effects of capturing and collaring on polar bears: findings from long-term research on the southern Beaufort Sea population. Wildl Res 41:311–322. doi: 10.1071/WR13225

Rosing-Asvid A, Born E, Kingsley M (2002) Age at sexual maturity of males and timing of the mating season of polar bears *(Ursus maritimus)* in Greenland. Polar Biol 25:878–883. doi: 10.1007/s00300-002-0430-7

Roulin A (2002) Why do lactating females nurse alien offspring? A review of hypotheses and empirical evidence. Anim Behav 63:201–208. doi: 10.1006/anbe.2001.1895

Rousset F (2008) GENEPOP'007: a complete re-implementation of the GENEPOP software for Windows and Linux. Mol Ecol Resour 8:103–106. doi: 10.1111/J. 1471-8286.2007.01931.X

Saunders BL (2005) The Mating System of Polar Bears in the Central Canadian Arctic. Queen’s University

Silva del Río N, Kirkpatrick BW, Fricke PM (2006) Observed frequency of monozygotic twinning in Holstein dairy cattle. Theriogenology 66:1292–1299. doi: http://dx.doi.org/10.1016/j.theriogenology.2006.04.013

Slate J, Visscher PM, MacGregor S, Stevens D, Tate ML, Pemberton JM (2002) A genome scan for quantitative trait loci in a wild population of red deer *(Cervus elaphus)*. Genetics 162: 1863–1873.

Spotte S (1982) The incidence of twins in pinnipeds. Canadian Journal of Zoology 60:2226–2233. doi: 10.1139/z82-285

Stirling I, Lunn NJ, Iacozza J (1999) Long-term trends in the population ecology of polar bears in western Hudson Bay in relation to climatic change. Arctic 52: 294–306.

Stirling I, Derocher AE (2012) Effects of climate warming on polar bears: a review of the evidence. Global Change Biol 18:2694–2706. doi: 10.1111/j.1365-2486.2012.02753.x

Szulkin M, Stopher KV, Pemberton JM, Reid JM (2013) Inbreeding avoidance, tolerance, or preference in animals? Trends Ecol Evol 28:205–211. doi: http://dx.doi.org/10.1016/j.tree.2012.10.016

Taylor M, Larsen T, Schweinsburg RE (1985) Observations of intraspecific aggression and cannibalism in polar bears *(Ursus maritimus)*. Arctic 38: 303–309.

Taylor MK, McLoughlin PD, Messier F (2008) Sex-selective harvesting of polar bears *Ursus maritimus*. Wildl Biol 14: 52–60.

Taylor RW, Boon AK, Dantzer B, RÉAle D, Humphries MM, Boutin S, Gorrell JC, Coltman DW, McAdam AG (2012) Low heritabilities, but genetic and maternal correlations between red squirrel behaviours. J Evol Biol 25:614–624. doi: 10.1111/j. 1420-9101.2012.02456.x

Thiemann GW, Iverson SJ, Stirling I (2008) Polar bear diets and arctic marine food webs: insights from fatty acid analysis. Ecol Monogr 78: 591–613.

Vibe C (1976) Preliminary Report on the Second Danish Polar Bear Expedition to North East Greenland, 1974. Proceedings of the Fifth Working Meeting of the Polar Bear Specialist Group. IUCN, Morges, Switzerland, pp 91–97

Villinger J, Waldman B (2012) Social discrimination by quantitative assessment of immunogenetic similarity.

Wang JL (2002) An estimator for pairwise relatedness using molecular markers. Genetics 160: 1203–1215.

Wang JL (2011) COANCESTRY: a program for simulating, estimating and analysing relatedness and inbreeding coefficients. Mol Ecol Resour 11:141–145. doi: 10.1111/J. 1755-0998.2010.02885.X

Weber DS, Van Coeverden De Groot PJ, Peacock E, Schrenzel MD, Perez DA, Thomas S Shelton JM, Else CK, Darby LL, Acosta L, Harris C, Youngblood J, Boag P, Desalle R (2013) Low MHC variation in the polar bear: implications in the face of Arctic warming? Anim Conserv 16:671–683. doi: 10.1111/acv.12045

Wilkinson GS (1992) Communal nursing in the evening bat, *Nycticeius humeralis*. Behav Ecol Sociobiol 31: 225–235.

Williams GC (1975) Sex and Evolution. Princeton University Press, Princeton, NJ

Zedrosser A, Støen O-G, Sæbø S, Swenson JE (2007) Should I stay or should I go? Natal dispersal in the brown bear. Anim Behav 74:369–376. doi: 10.1016/j.anbehav.2006.09.015

Zeyl E, Aars J, Ehrich D, Bachmann L, Wiig Ø (2009a) The mating system of polar bears: a genetic approach. Can J Zool 87:1195–1209. doi: 10.1139/Z09-107

Zeyl E, Aars J, Ehrich D, Wiig Ø (2009b) Families in space: relatedness in the Barents Sea population of polar bears *(Ursus maritimus)*. Mol Ecol 18:735–749. doi: 10.1111/J.1365-294x.2008.04049.X

